# Exploring the diversity of anti-defense systems across prokaryotes, phages, and mobile genetic elements

**DOI:** 10.1101/2024.08.21.608784

**Authors:** Florian Tesson, Erin Huiting, Linlin Wei, Jie Ren, Matthew Johnson, Rémi Planel, Jean Cury, Yue Feng, Joseph Bondy-Denomy, Aude Bernheim

**Author notes:** The authors wish it to be known that, in their opinion, the first 2 authors should be regarded as joint First Authors.

## Abstract

The co-evolution of prokaryotes, phages, and mobile genetic elements (MGEs) over the past billions of years has driven the emergence and diversification of defense and anti-defense systems alike. Anti-defense proteins have diverse functional domains, sequences, and are typically small, creating a challenge to detect anti-defense homologs across the prokaryotic genomes. To date, no tools comprehensively annotate anti-defense proteins within a desired genome or MGE. Here, we developed “AntiDefenseFinder” – a free open-source tool and web service that detects 156 anti-defense systems (of one or more proteins) in any genomic sequence. Using this dataset, we identified 47,981 anti-defense systems distributed across prokaryotes, phage, and MGEs. We found that some genes co-localize in “anti-defense islands”, including *E. coli* T4 and Lambda phages, although many are standalone. Out of the 112 systems detected in bacteria, 100 systems localize only or preferentially in prophages, plasmids, phage satellites, integrons, and integrative and conjugative elements. However, over 80% of anti-Pycsar protein 1 (Apyc1) resides in non-mobile regions of bacteria. Evolutionary and functional analyses revealed that Apyc1 likely originated in bacteria to regulate cNMP signaling, but was co-opted multiple times by phages to overcome cNMP-utilizing defenses. With the AntiDefenseFinder tool, we hope to facilitate the identification of the full repertoire of anti-defense systems in MGEs, the discovery of new protein functions, and a deeper understanding of host-pathogen arms race.

## INTRODUCTION

In the last several years, there have been over a hundred newly identified systems in prokaryotes that defend against phages^1^. Several studies have revealed mechanistic diversity of defense systems, spanning nucleic acid^2–9^ or metabolite^10–16^ depletion, signaling molecule cascades^17–20^ membrane disruption^21–23^ and many more^24–26.^ To counteract these systems, phages evolved a diversity of anti-defense systems that directly inhibit individual defense proteins^7,10,26–29^ or signaling molecules^29–36^ or indirectly inhibit these systems through reversal of defense function^37^.

To date, the most well-studied anti-defense strategies are anti-Restriction-Modification (RM) and anti-CRISPR proteins that provide protection against nucleic acid targeting systems. These proteins have been extensively studied in phage, prophages^38,39^, plasmids^40^, and conjugative elements^39^. In certain cases, MGE-encoding anti-CRISPR proteins that inhibit Type III CRISPR-Cas systems^41^ have been co-opted by the bacterial host to regulate the Type III CRISPR-Cas activity^42^. Beyond inhibitors of CRISPR-Cas and RM, the distribution and localization of other anti-defense systems remains vastly understudied. The main challenge in identifying anti-defense proteins is due to the vast diversity of the functional domains and the often small protein size (i.e. 80% of anti-defense proteins are smaller than 200 amino acids), a bottleneck for both sequence and structure-based detection. creating

To address this, we built upon the established DefenseFinder^1,43,44^ search tool and web service to detect all known anti-defense systems in prokaryotic and phage genomes. Since the discovery of the first anti-restriction protein^45^ there have been at least 180 proteins identified to inhibit prokaryotic defense systems. A pre-computed database of 41 experimentally validated anti-defense systems (dbAPIS) was published that identified 4,428 homologs of anti-defense systems in phages^46^. Our newly developed AntiDefenseFinder tool can detect 156 anti-defense systems (some systems are composed of multiple proteins). When applied to the RefSeq database of 21,855 prokaryotic complete genomes and from the GenBank database of 13,487 phage sequences, it detects 41,972 and 6,009 anti-defense systems in prokaryotic and phage genomes, respectively. Alongside this comprehensive dataset, the search tool is available on a freely accessible web service and via command line, which we hope will facilitate the identification of anti-defense genes within any DNA or protein sequences.

We found that most anti-defense systems are variable in frequency and distribution across prokaryotic species. We observed several instances of anti-defense genes co-localizing into “anti-defense islands”, including the model *E. coli* T4 and Lambda phages. In some cases, these anti-defense islands contain only anti-defense genes from a single family, such as anti-CRISPRs, anti-Gabija, or anti-Thoeris. However, many anti-defense genes tend to be encoded alone across a combination of prophages, plasmids, phage satellites, integrons, and integrative and conjugative elements. We also identified that NAD+ reconstitution pathway 1 and 2 (NARP1/2) and anti-Pycsar (Apyc1) genes are enriched in non-MGE sequences within the bacterial chromosome. Based on our evolutionary and functional analyses, we predict that Apyc1 homologs are common in prokaryotic genomes to regulate housekeeping signals, such as cAMP, and this cNMP-cleaving protein was co-opted by phages to counteract defense systems using cCMP and cUMP. This newfound understanding of Apyc1 sets a precedent for in-depth, quantitative bioinformatic evaluations of anti-defense systems to uncover further insights into the ongoing host-pathogen arms race.

## MATERIALS AND METHODS

### Databases used in the study

Two databases were utilized in this study. First, we used the RefSeq complete genome database for bacteria and archaea, which was downloaded in July 2022 and contains 21,855 genomes. For phage genomes, we utilized the GenBank database, which was downloaded in December 2023 and includes 13,487 genomes.

### Protein sequence models

All experimentally validated protein sequences were retrieved from the literature (Table S4). All proteins were blasted using BLASTp against the NCBI non-redundant database with an E-value threshold of 1e-5. The resulting hits were then compared to the original protein sequence to ensure a minimum of 30% identity. Additionally, a coverage threshold was applied: 80% of coverage of the original protein and 70% of coverage of the hit (i.e. the 70% of the hit protein corresponds to the original protein). All conserved hits were then clustered at 95% identity and 95% coverage using Mmseqs2^47^ v13.45111 easy-cluster. If the number of representative sequences was higher than 200, the sequences were clustered at 80% coverage and 80% identity. All representative sequences were then aligned using mafft^48^ v7.505 (default settings) and hmm profiles were built using hmmbuild (HMMER^49^ v3.3.2).

### Mobile genetic element and defense system detection

RefSeq annotation was used to determine if a given replicon was a plasmid. Prophages were detected using Virsorter2^50^ v2.2.3. An anti-defense system was classified as inside a prophage if it was present in the boundaries of the prophage (+/-2kb). Satellites were detected using SatelliteFinder v0.9.1. An anti-defense system was classified as inside a satellite if it was present in the boundaries of the prophage (+/-2kb). Integrons were detected using IntegronFinder^51^ v2.0.2. An anti-defense system was classified as inside an integron if the protein was detected as part of an integron cassette by IntegronFinder ICE were detected using CONJScan Macsyfinder models^52^ v2.0.1 to detect conjugative systems on chromosomal replicon (not annotated as plasmid). An anti-defense system was classified as inside an ICE if it was present between the extremities of the detected proteins +/-10kb. All integrases were detected using 108 PFAM^53^ with the PFAM description containing “Transposase”, “Recombinase”, “Integrase” and “Resolvase” using GA thresholds with Hmmsearch (HMMER^49^ v3.3.2).

### First detection of anti-defense system and threshold choice

All profile HMMs detection was done using Hmmsearch (HMMER^49^ v3.3.2) on both the prokaryotic RefSeq database and Genbank phage database with GA cut threshold at 20 and profile coverage of 40%. All hits were then classified between 4 categories based on their localization: Phage (Genbank database), Plasmid, prophage or Other. All GA (hit score) thresholds were manually chosen. Those thresholds were defined using three main factors: hit score, coverage distribution and hit localization in the genome. These criteria were combined in a single graph illustrated in Figure 1B and available for genes with more than 1,000 hits Figure S1 and for all profiles on GitHub (https://github.com/mdmparis/antidefensefinder_2024) and on Figshare under the DOI: 10.6084/m9.figshare.26526487.

**Figure 1.**
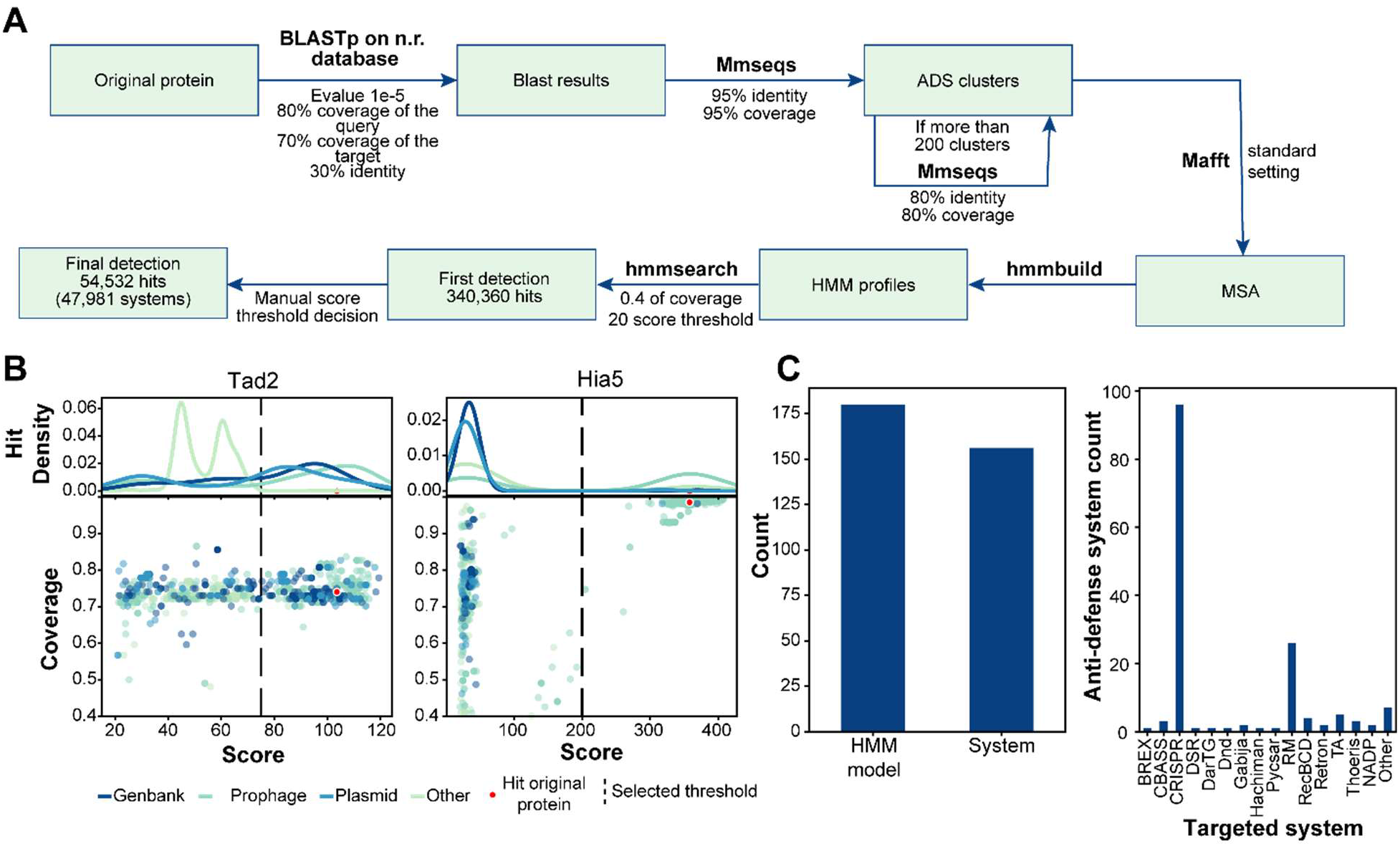
AntiDefenseFinder is a tool to systematically detect known inhibitors of prokaryotic defense systems. **(A)** Pipeline of creating HMM profiles for AntiDefenseFinder. **(B)** Filtering of positive hits based on selected threshold and protein sequence coverage (> 40%). The selection threshold for each anti-defense protein was manually analyzed and chosen based on the distribution of hits relative to the originally discovered protein. **(C)** Total number of HMM models developed relative to the total number of anti-defense proteins, and total number of anti-defense systems detected across prokaryote and phage sequences that inhibit a specific type or family of defense systems.

### Anti-defense system and defense system detection

Anti-defense system and defense systems were detected using defense-finder v1.3.0 with the argument --antidefense on the two databases.

### Apyc1 phylogenetic tree

All Apyc1 homologs detected by AntiDefenseFinder were retrieved. Bacterial homologs were clustered together at 80% identity and 80% coverage with Mmseqs2^47^ v13.45111. Phage homologs were clustered with Mmseqs2 at 95% identity and 95% coverage. All representative sequences were used for the alignment. 18 sequences of Metallo Beta Lactamase (MβL) fold protein known to be antimicrobial resistance genes were used as an outgroup of the tree. The alignment used for the tree construction was made using muscle^54^ v5.1 with the -super5 option. The alignment was trimmed using clipkit^55^ v1.3.0 in smart gap mode. The tree was built using IQ-TREE^56^ v2.2.3 with models finder and 2000 ultrafast bootstrap.

### Apyc1 multiple sequence alignment

Apyc1 protein sequences in Figure 4 were aligned using EMBL-EBI MUSCLE and then visualized using Jalview v2.11.3.3. These Apyc1 sequences included: *Thalassospira* WP_223304948.1 (THSP027), *Archangium violaceum* WP_204220610.1 (ARVI001), *Bacillus* phage SBSphiJ (Hobbs et al. 2022), *Paenibacillus sp. J14* WP_028539944.1 (PASP001), *ohnella* WP_174887610.1 (COSP018), *Legionella sp. MW5194* WP_203455517.1 (LESP016), *Synechocystis* WP_010871596.1 (SYSP007), *Staphylococcus* phage Madawaska QQO92874.1 (MW349129), *Caldicellulosiruptor bescii* WP_041727399.1 (CABE001).

### Apyc1 protein structure predictions

Apyc1 protein sequences in Figure 4 (listed above) were predicted using AlphaFold2 ColabFold^57^ v1.5.5. Structural comparison of the Apyc1 proteins was performed using the super function in Pymol v2.1.

### Apyc1 protein purification

The *apyc1* genes were synthesized and cloned into pET28a vectors in which the expressed protein contains an N-terminal His6 tag. All the proteins were expressed in *E. coli* strain BL21(DE3) in lysogeny broth (LB) medium. After growth at 37°C, the cells were induced by 0.2 mM isopropyl-β-d-thiogalactopyranoside (IPTG) when the cell density reached an optical density at 600 nm of 0.8. After growth at 18°C for 12 h, the cells were harvested, resuspended in lysis buffer (50 mM Tris–HCl pH 8.0, 300 mM NaCl, 30 mM imidazole and 1 mM PMSF) and lysed by sonication. The cell lysate was centrifuged at 20,000 g for 50 min at 4°C to remove cell debris. The supernatant was applied onto a self-packaged Ni-affinity column (2 mL Ni-NTA, Genscript) and contaminant proteins were removed with washing buffer (50 mM Tris-HCl pH 8.0, 300 mM NaCl, 30 mM imidazole). Then the protein was eluted with an elution buffer (50 mM Tris pH 8.0, 300 mM NaCl, 300 mM imidazole). The protein eluent was concentrated and further purified using a Superdex-200 increase 10/300 GL (Cytiva) column equilibrated with a buffer containing 10 mM Tris-HCl pH 8.0, 200 mM NaCl and 5 mM DTT. For the LESP016-Apyc1 and MW349129-Apyc1, buffers contained 500 mM NaCl along with an additional 5% glycerol throughout the purification process.

### Apyc1 *in vitro* cleavage assays

Reactions of the assay consisted of 50 mM Tris-HCl pH 7.5, 100 mM KCl, 1 mM MgCl_2_, 1 mM DTT, 100 µM cNMP and 1 µM recombinant protein in a 100 μL volume. The reaction mix was incubated at 37°C for 20 min and then filtered using a 3-kDa cutoff filter (Millipore) at 4°C. Filtered nucleotide products were analyzed using a C18 column (Agilent ZORBAX Bonus-RP 4.6 × 150 mm) heated to 30°C and run at 1 ml/min in a buffer of 50 mM NaH_2_PO_4_ adjusted to pH 6.8, supplemented with 3% acetonitrile and 0.1% trifluoroacetic acid. Raw data provided in Figure S6.

### Apyc1 enzymatic kinetics assays

The kinetic experiments were conducted at 37°C with a total reaction volume of 100 μL, in a buffer containing 50 mM Tris-HCl pH 7.5, 100 mM KCl, 1 mM DTT, and 1 mM MgCl_2_. Reactions were initiated by adding protein and proceeded for 20 seconds, then they were terminated with 0.1 M NaOH. Subsequently, the reaction samples were placed into the HPLC autosampler. Each reaction mix was analyzed using the C18 column under the above conditions. The area of the substrate peak at 254 nm was integrated to determine the substrate consumption at each substrate concentration. The data were converted into reaction rates and plotted against substrate concentrations. Curve fitting and kinetics parameter determination were performed using the Origin software. Raw data provided in Figure S7.

## RESULTS

### Anti-DefenseFinder: A search tool to detect known inhibitors of prokaryotic defense systems

To systematically detect anti-defense systems, we developed an AntiDefenseFinder option, to DefenseFinder^1,43,44^, a program that already detects defense systems. We first conducted a comprehensive literature review of all known anti-defense proteins and retrieved experimentally validated sequences of 180 anti-defense genes. DefenseFinder relies on Hidden Markov Model (HMM) profiles for sensitive homology search. We thus needed to build one HMM profile per anti-defense protein. To automate the creation of the HMM profiles, we started with a homology search using BLASTp on the RefSeq non-redundant database to capture sequence diversity (Figure 1A). BLASTp results were filtered using a minimal coverage and sequence identity. Next, the sequences were clustered to reduce the weight of closely related homologs (for example, *Escherichia coli* proteins) in the multiple sequence alignment. Cluster representatives were then aligned and a HMM profile was constructed for 156 families of anti-defense systems because several proteins may be part of the same family (e.g. ADPS and Namat) or can be found in several families (e.g. Anti-CRISPR associated (Aca) proteins).

We performed initial detections on two databases: RefSeq prokaryotic complete genomes and GenBank phage genomes (See Methods). Using a low threshold (GA: 20 and coverage > 40%), we identified 340,360 hits (Table S1). These hits were used to refine each HMM profile’s GA threshold (hit score) based on the distribution of both hit scores and profile coverage (Figure S1). As anti-defense genes are often encoded inside mobile genetic elements (MGEs), we also took into consideration the localization of hits within genomes or MGEs to further define a true positive hit (Figure 1B). Hits within MGEs (e.g., plasmids, prophages, or phage databases) were considered more likely to be true positives. This approach allowed us to manually set a threshold for each profile (Figure S1). Overall, AntiDefenseFinder detects 156 anti-defense systems with HMM profiles encompassing 180 proteins (Figure 1C). The majority of known anti-defense systems are anti-CRISPR (n=96) and anti-Restriction-Modification (anti-RM) (n=26), but AntiDefenseFinder also identifies a variety of other anti-defense systems that target the expanding diversity of prokaryotic defense systems. AntiDefenseFinder is now integrated into DefenseFinder version v1.3.0 available in command line and as a web service. It can be executed alongside DefenseFinder (--antidefensefinder) or using only AntiDefenseFinder models (-- antidefensefinder-only).

### Anti-defense systems are variably distributed across genomes and genetic elements

We initially sought a comprehensive view of anti-defense system distribution across prokaryotes and phages. To do so, we applied AntiDefenseFinder to a database of 21,855 prokaryotic and 13,487 phage genomes and detected a total of 47,981 anti-defense systems. In bacteria, 41,946 total anti-defense systems were identified and were significantly enriched in bacteria of the genera *Escherichia* (12,544 total, ∼25%), *Klebsiella* (9,108), *Staphylococcus* (1,781), *Enterococcus* (1,242), *Pseudomonas* (579), and *Bacillus* (320) (Figure 2A and 2C, Table S2). Anti-RM and anti-CRISPR (Acr) are the most abundant anti-defense systems in bacteria with a total count of 22,708 and 6,880, respectively (Figure 2A and 2C). In *Escherichia*, anti-RM systems are the most abundant and are notably enriched with 3,132 instances of ArdB/KlcA. This may have occurred because ArdB was discovered in *Escherichia coli* in 1993^45^, and has henceforth been studied in-depth in the same bacteria host. In *Pseudomonas*, Acr systems are the most abundant and are enriched with Type I and II CRISPR-Cas Acr proteins, but most notably 114 instances of AcrIF3. This again may be due to the discovery of Acrs in *Pseudomonas aeruginosa*^38,58^. Apart from anti-RM and Acrs, 43% (10/23) of anti-defense systems with greater than 10 instances are only detected in the phylogenetic order where the system was originally discovered. Otherwise, anti-defense systems are variable between bacterial species. For instance, in *Klebsiella*, the anti-Pycsar protein 1 (Apyc1) is the most abundant with 1,233 homologs detected, and in *Acinetobacter*, the newly identified NAD+ reconstitutions pathway 1 (NARP1) is the most abundant with 469 occurrences. In the 383 genomes of archaea, only 26 anti-defense systems were detected and 65% of those systems were Acrs (AcrIII1 n=7, AcrIIA26 n=7, AcrIA1 n=3). Only five anti-defense systems detected were not Anti-RM or Acrs.

**Figure 2:**
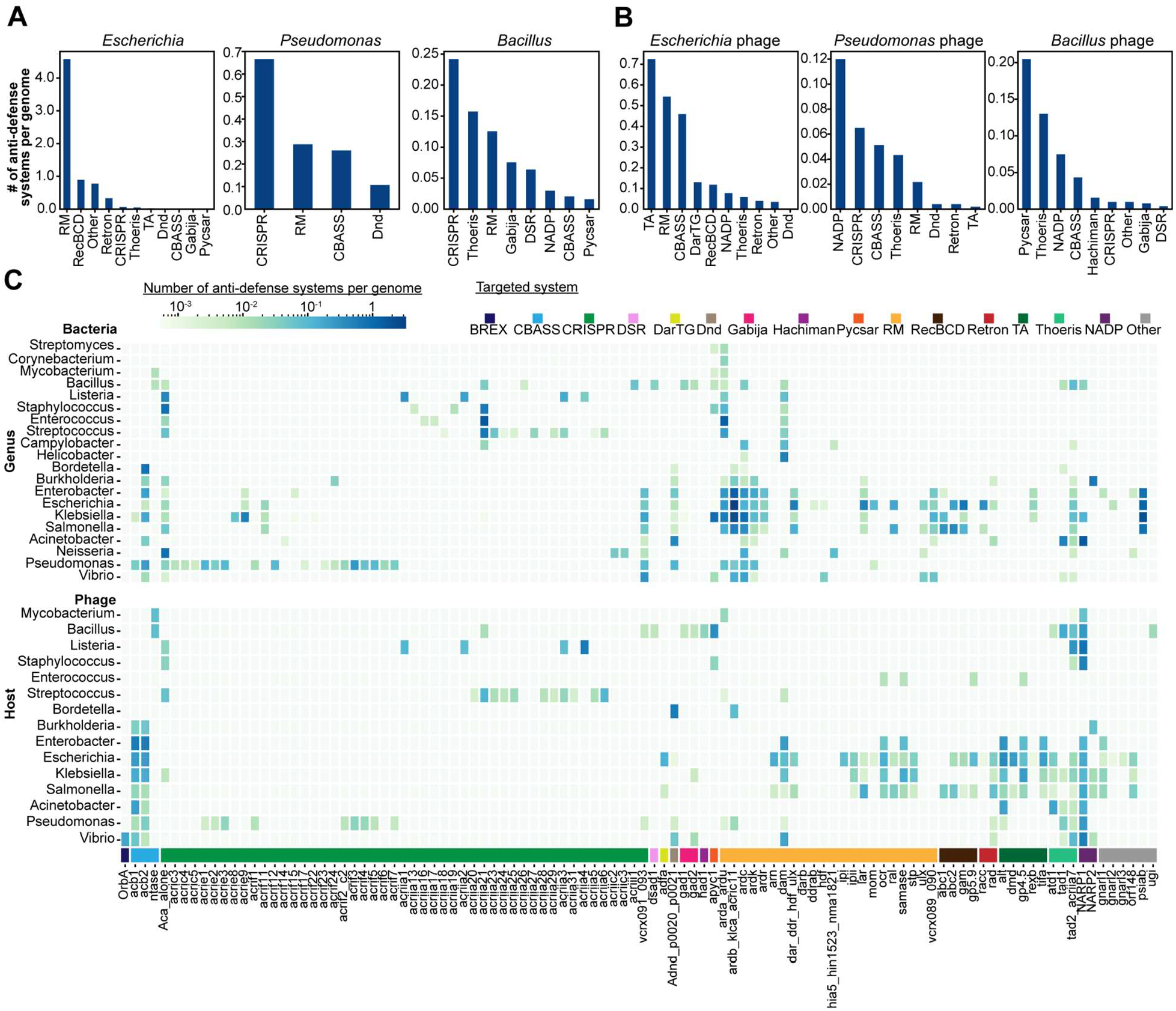
Anti-defense system distribution across different bacterial genera and phage host. Total anti-defense systems found across **(A)** bacteria or **(B)** phages in *Escherichia, Pseudomonas*, or *Bacillus* genera. **(C)** Average number of ADS found per genome organized by genus.

In phages, 6,009 total anti-defense systems were identified and enriched in phage that infect the genera *Escherichia* (2,124 total, ∼35%), *Klebsiella* (453), *Vibrio* (321), *Salmonella* (299), *Pseudomonas* (158), *Bacillus* (254) (Figure 2B and 2C; S2, Table S3). Similarly to bacteria, anti-RM is the most abundant anti-defense system. We suspect this is due to a bias in the available genomic sequences and the early discovery and the prevalence of RM in bacteria. Aside from anti-RM, 56% (14/25) of anti-defense systems with greater than 10 instances are only detected in the phylogenetic order where they were discovered (e.g. ArdB in *Enterobacterales*, AcrIIA1 in *Bacillales* or AcrIIA23 in *Lactobacillales*). For example, anti-CBASS protein 1 (Acb1) and Acb2 are enriched in phage genomes infecting eight related genera (Figure 2C). There are also instances when anti-defense systems are only found in phage (e.g. Had1, Ocr, etc.) or only in bacterial genomes (e.g. AcrIIA13, PsiAB, etc.). In any case, many anti-defense systems are very rare and present in less than 1% of prokaryotic and phage genomes. Overall, these results demonstrate that anti-defense system distribution is variable across distinct prokaryotic and phage genomes with a bias towards model organisms where they were originally identified in. This suggests that discovery of anti-defense in new species is important toward a better understanding of the anti-defense diversity.

We then set out to understand how anti-defense systems are localized across the prokaryotic pan-genome and mobile genetic elements (MGEs). Anti-CRISPR (Acr) genes typically co-localize or become enriched in genomic loci of prophages^38,39^ and anti-RM, anti-SOS, and Acrs can co-localize on the leading strand of conjugative plasmids^40^, which has been collectively referred to as “anti-defense islands”. We therefore evaluated whether this observation could extend to other anti-defense systems and observed that anti-defense systems co-localize within 10kb of one another in 31.7% and 32.9% of bacterial and phage genomes, respectively, with 17.8% of systems co-localizing within 1kb in phage genomes. The well-studied *E. coli* T4 phage has at least three independent instances of anti-defense genes co-localizing together in an anti-defense island while the *E. coli* phage Lambda has one instance (Figure 3A). Other co-localization of anti-defense systems occurs in phages from the BASEL collection, such as Bas31 and Bas35 (Figure S3). In all these cases, these anti-defense islands include genes that have been shown to inhibit distinct bacterial defense systems. In other phages, anti-defense genes that target the same bacterial defense system co-localize in the genome, such as Acrs, anti-Gabija (Gad), and anti-Thoeris (Tad) genes across *Pseudomonas, Bacillus*, and *Blautia* bacterial genomes, respectively (Figure 3B). We anticipate that more anti-defense islands are present in MGEs due to the increasing identification and diversity of anti-defense systems. That withstanding, these results demonstrate that ∼66% of all known anti-defense genes do not localize in the same genetic loci, but rather encoded alone (Figure S2). As an example, applying Anti-DefenseFinder to the well-studied *P. aeruginosa* model jumbo phage phiKZ revealed only one anti-defense gene, Tad1, despite encoding dozens of small genes of unknown function. These collective results reflect a need for discovering new anti-defense genes.

**Figure 3:**
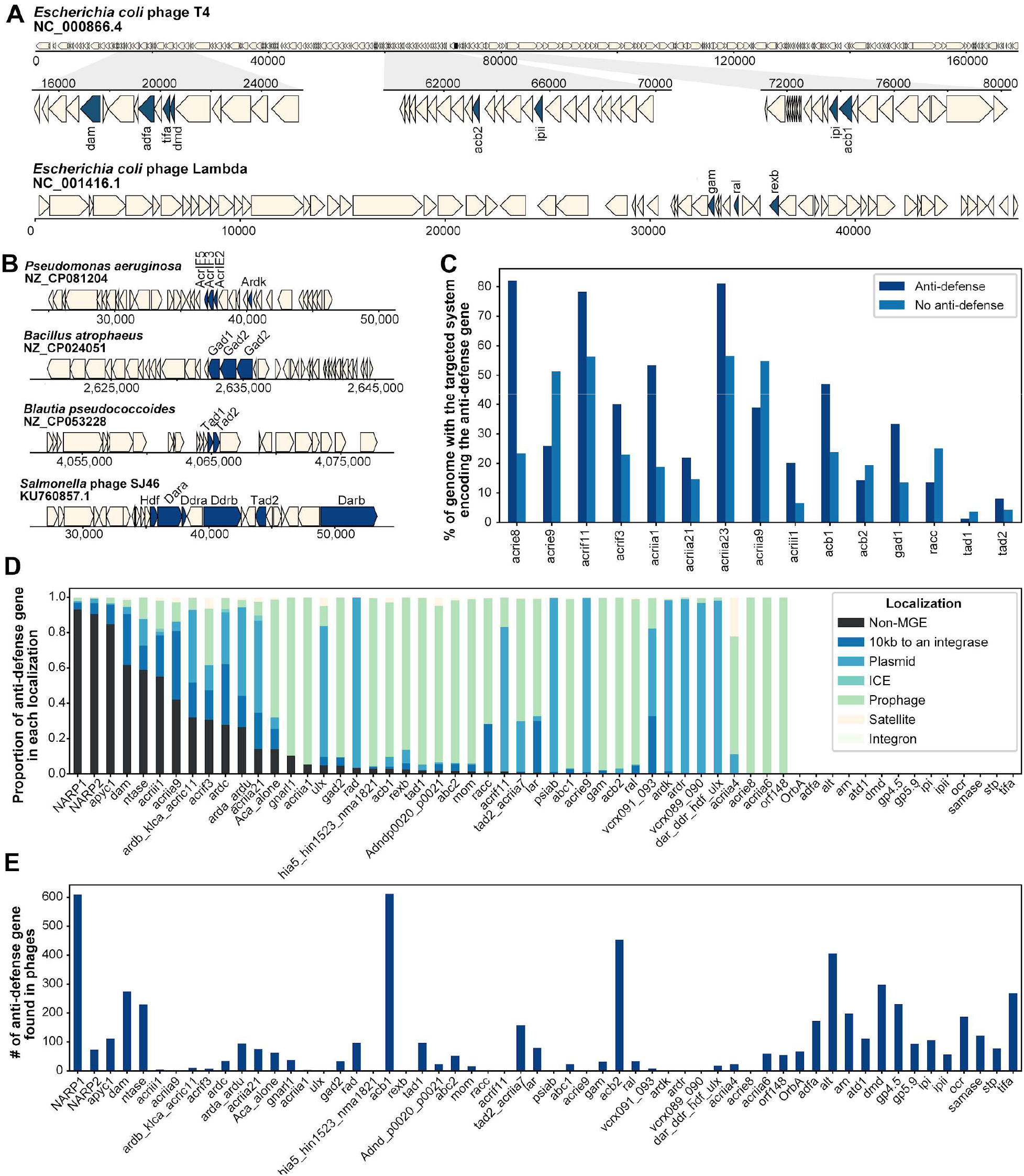
Localization of anti-defense systems in genomes and MGEs. **(A)** Examples of anti-defense genes co-localized in an anti-defense island within the well-studied *E. coli* phages T4 and Lambda, and **(B)** diverse bacterial and phage genomes. **(C)** Conditional percentage of genome encoding the targeted system when the anti-defense system is encoded or not. **(D)** Relative proportion of a single anti-defense genes localized in distinct genomic localizations, including: satellites, prophages, integrative conjugative elements (ICE), plasmids, nearby integrases, and not in mobile genetic elements (MGEs) like bacterial chromosomes. **(E)** Total number of anti-defense genes localized in phage genomes.

We next evaluated whether anti-defense systems are encoded in the same genome as the defense system they were originally identified to inhibit. We found that most anti-defense genes do not appear to co-occur in the same genome as its targeted defense system (Figure 3C). However, AcrIE8 is a unique example that is often encoded in genomes that also encode Type I CRISPR-Cas (Figure 3C). Nearly all instances of AcrIE8 are encoded on prophages (Figure 3D), with previous work suggesting that anti-CRISPRs are expressed to neutralize CRISPR and prevent self-targeting^38,39^. Some other Acr genes (i.e. *acrIF11, acrIIA1*, and *acrIIA23*) are often found in the same genome with the CRISPR-Cas system they inhibit. Expanding upon this analysis revealed that many anti-defense genes are encoded in MGEs, including satellites, prophages, integrative conjugative elements (ICE), plasmids and nearby integrases, and fewer anti-defense genes are encoded in non-mobile regions (Figure 3D). In many cases, >80% of instances of the detected anti-defense gene are encoded within a single type of MGE (Figure 3D), suggesting that the inhibited defense system may predominantly target that type of MGE.

We hypothesized that identifying anti-defense genes encoded on a distinct type of MGE would reveal an unexpected target of the defense system. However, our findings generally align with the known defense system mechanism. For example, we observed that anti-RM and Acr genes are encoded on diverse types of MGEs (Figure 3D), and it is known that RM and CRISPR-Cas systems target various MGEs^59,60^. By comparison, anti-CBASS (Acb) and Tad genes are primarily encoded on phages and prophages (Figure 3D and 3E). To date, both CBASS and Thoeris have been demonstrated to only target phages^17,18,32^. Other anti-defense genes are only encoded in lytic phages, including Ocr (anti-RM; *Teseptimavirus* and *Kayfunavirus* phage)^61^, Had1 (anti-Hachiman; *Bastillevirinae* phage)^62^, Atd1 (anti-TIR; *Phapecoctavirus, Justusliebigvirus, Lazarusvirus* phage)^36^, and AdfA (anti-DarTG; *Tequatrovirus, Mosigvirus* phage)^26^ (Figure 3E). In several of these cases, the cognate defense system has been demonstrated to solely target phages. Surprisingly, our final analyses demonstrated that a limited number of anti-defense genes are enriched in non-mobile regions of the bacterial genome, such as anti-Pycsar (Apyc1)^31^, NAD+ reconstitution pathway 1 and 2 (NARP1/2)^37^, and NTases (anti-CBASS)^36^ (Figure 3D), suggesting a non-defenses function for these proteins. We further investigate bacterial and phage Apyc1 below.

### Anti-Pycsar gene is common in bacterial chromosome and co-opted by phages

The pyrimidine cyclase system for anti-phage resistance (Pycsar) uses cCMP or cUMP signaling molecules to activate a downstream effector that acts on the bacterial host and induces premature cell death, limiting phage replication^19^. In response, phage evolved anti-Pycsar protein 1 (Apyc1) that counteracts this system through cleavage of cyclic mononucleotides (cCMP, cUMP, cGMP, cAMP)^31^. This study also identified 107 Apyc1 homologs in distinct phages and bacterial chromosomes in two predominant *Bacillus* and *Staphylococcus* clades and then 10 homologs were experimentally validated to cleave cCMP and cUMP^31^. Using the AntiDefenseFinder tool, we detected 2,301 total instances of Apyc1 with 80.7% encoded in the bacterial chromosome outside of an obvious MGE (Figure 3C). To determine the evolutionary history of Apyc1 homologs, we built a phylogenetic tree of bacterial and phage Apyc1 and used an antimicrobial resistance MβL-fold protein as an outgroup to root the tree. We observed three independent monophyletic clades of phage Apyc1 branching in bacterial Apyc1 (Figure 4A), suggesting that bacterial Apyc1 represents the ancestral form that phage likely acquired Apyc1 from a bacterial host in multiple events. Upon further investigation, we observed that bacterial Apyc1 is encoded in genomes that also include Pycsar, CBASS, and occasionally, Apyc1 is adjacent to a cyclase with no obvious effector nearby (Figure S4).

**Figure 4:**
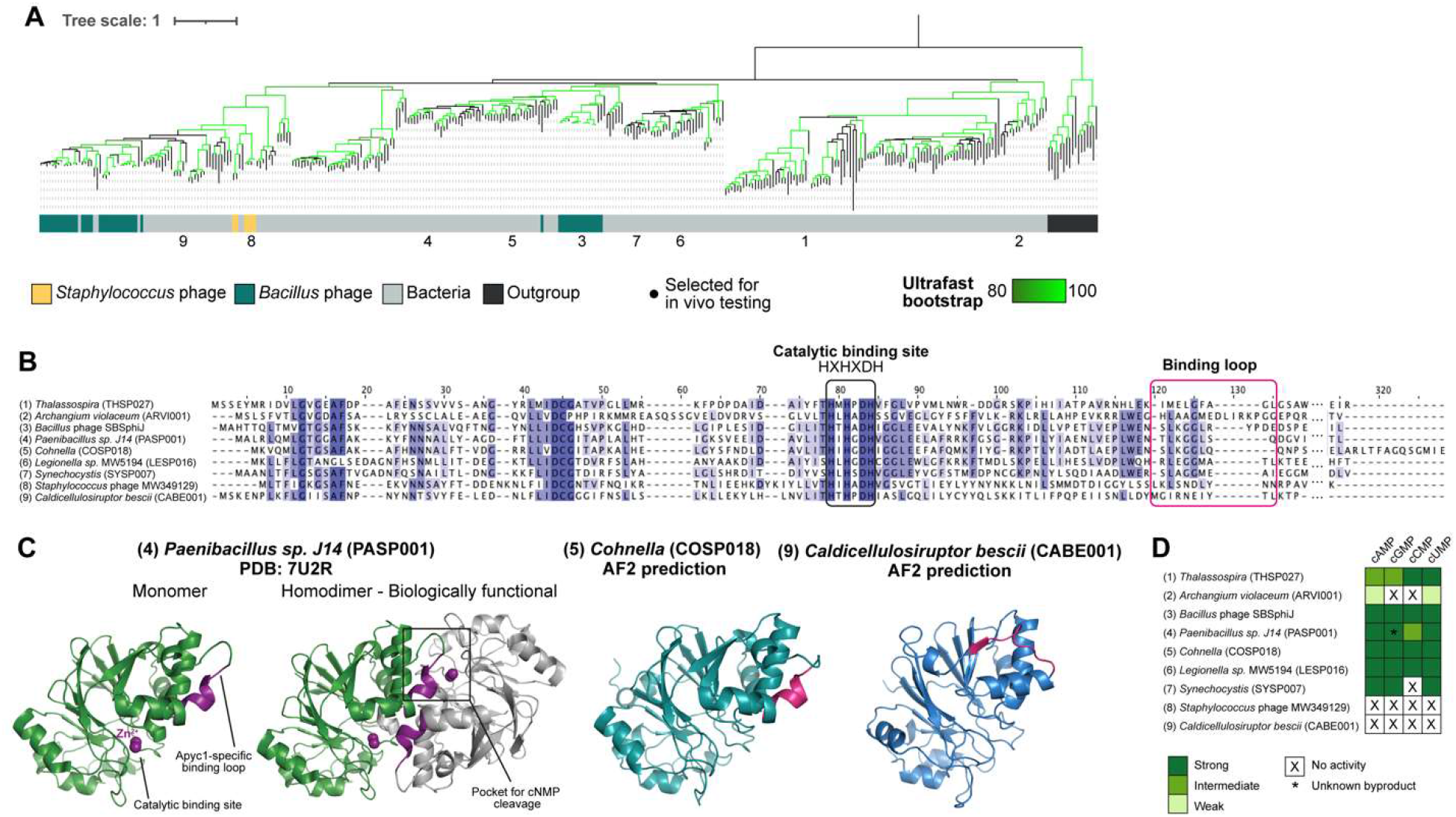
Anti-Pycsar (Apyc1) is abundant and functionally conserved across bacteria. (A) Phylogenetic tree of SBSphiJ Apyc1 and >350 homologs from bacteria and phage. Colors represent bacterial genus, highlighting the most abundant *Bacillus* and *Staphylococcus*. Black circles indicate Apyc1 homologs tested for *in vitro* cleavage of cyclic mononucleotides (cNMPs). (B) Multiple sequence alignment of Apyc1 homologs (see Figure S4 for full alignment). Residues that are >80 % conserved, >60 % conserved and >40% conserved are shaded in dark purple, light purple, and light gray, respectively. Residues involved in catalysis and binding are circled in black and pink, respectively. **(C)** Structures of *Paenibacillus sp. J14* (PASP001), *Cohnella* (COSP018), and *Caldicellulosiruptor bescii* (CABE001) Apyc1 homologs. PASP001 was experimentally solved and deposited on the RCSB Protein Data Bank (PBD: 7U2R), and COSP018 and CABE011 were computationally predicted using AlphaFold2 (AF2). Zn^2+^ ions that coordinate cNMP cleavage in the catalytic binding site, as well as the Apyc1-specific loop that extends into the cNMP catalytic binding site, are labeled and highlighted in pink. **(D)** Summary of the *in vitro* cleavage assay data (n=3).

To determine whether bacterial Apyc1 is functional, we initially examined the sequence and structure of evolutionarily diverged Apyc1 homologs in bacteria and phage. We observed that the Apyc1 sequences all retain the catalytic site, but exhibit diversity in the nucleotide binding loop (Figure 4B), which is proposed to extend into the nucleotide-binding pocket and stabilize the small cyclic mononucleotide substrates^31^. For the *Paenibacillus sp. J14* Apyc1 homolog (PASP001), the structure was previously solved and demonstrated that the loop from one monomer interacts with the catalytic binding pocket of the opposing monomer and subsequently enables cCMP hydrolysis^31^ (Figure 4). For the bacterial homologs we examined, such as *Cohnella* (COSP018), the nucleotide binding loop is intact and overlays well with PASP001 loop (Figure 4C and Figure S5), suggesting that it also retains the nucleotide cleavage function. Some bacterial homologs like *Caldicellulosiruptor bescii* (CABE001) exhibit a shortened loop (Figure 4C) while others exhibit a lengthened loop (Figure S5), and in turn, may not effectively interact with the catalytic binding pocket.

To examine the function of these Apyc1 homologs, we performed *in vitro* cleavage assays and observed that bacterial Apyc1 homologs with structurally conserved nucleotide binding loops were able to strongly cleave cAMP, cGMP, cCMP, and cUMP signals (Figure 4D and S6). The PASP011 and SBSphiJ Apyc1 homologs examined in Hobbs et al. (2022) also demonstrated cleavage of all cNMP signals. By comparison, CABE001, *Archangium violaceum* (ARVI001), and *Staphylococcus* phage (MW349129) homologs with shortened or lengthened Apyc1-specific nucleotide binding loops showed weak or no cleavage of cNMPs (Figure 4D). These data suggest that the bacterial Apyc1 function is degradation of cNMPs, which was then co-opted by phages. Finally, we investigated whether the phage versions of the enzyme displayed faster turnover compared to the host version. To do so, we examined enzymatic kinetics from Apyc1 homologs in the *Bacillales* order – bacterial PASP011 and COSP018 and phage SBSphiJ Apyc1 – with the Pycsar signals cCMP and cUMP. We observed that the bacterial COSP0018 Apyc1 homolog cleaves cCMP and cUMP with nearly identical kinetics compared to phage Apyc1 while the bacterial PASP011 Apyc1 demonstrated ∼6-fold and ∼2-fold slower kinetics with cCMP and cUMP, respectively (Figure S7). In addition, we observed that all Apyc1 homologs exhibit high K_cat_ values (275-1,581 s^-1^) (Figure S7). These findings suggest that bacterial and phage Apyc1 have generally similar enzyme kinetics without specialization or adaptation by the phage homologs. Altogether, we conclude that the Apyc1 family functions in rapid cleavage of cNMPs that are likely utilized in both regulatory and defense systems.

## DISCUSSION

We developed the AntiDefenseFinder tool and web-service (https://defensefinder.mdmlab.fr) that detects all known anti-defense systems across prokaryotic and phage genomes, as well as mobile genetic elements (MGEs). In doing so, we provided a quantitative overview of 156 anti-defense systems families and 47,981 homologs that span a diversity of bacterial genera, genomic localizations, and functional strategies. A recently developed pre-computed database, dbAPIS, detects 41 anti-defense systems and 4,428 total homologs encoded in phage genomes^46^. We hope that the free and open-source, searchable nature of AntiDefenseFinder will enable the field to identify the full repertoire of anti-defense systems, especially in understudied MGEs. AntiDefenseFinder is also easily adaptable to add new anti-defense systems given that we built upon the pre-existing framework of the DefenseFinder tool^1,43,44^. Over time, we will continue building new profiles of anti-defense systems.

Many gaps of knowledge remain regarding anti-defense systems, such as species diversity and anti-defense island abundance. Although we observed anti-defense genes widespread across many distinct prokaryotic species, there is biased enrichment in *Escherichia* (14,668 detected) and related species, likely because these model organisms were used to discover the first instance of the anti-defense gene. In both bacterial and phage genomes, we also observed that over 30% of detectable anti-defense genes co-localize within 10kb of one another in bacteria and 17.8% within 1kb in bacteriophages, which is a defining feature of anti-defense islands. The model *E. coli* T4 phage notably encoded three independent instances of anti-defense islands; however, many bacteria and phage still lack these islands. Conversely, there are over 60% of anti-defense genes that are standalone. It is possible that applying a “guilt-by-association” analysis may reveal entirely new anti-defense genes as it did with anti-CRISPRs (Acrs)^39,63^. We anticipate that an abundance of anti-defense systems await discovery in prokaryotic host species that currently lack known anti-defense genes or islands.

Challenges remain in the detection of distantly related anti-defense proteins due to their small size and vast sequence divergence. In some cases, the functional domains of enzymatic proteins are widely conserved, like the phosphodiesterase domain of anti-CBASS protein 1 (Acb1)^31^. Enzymatic domains have been found to retain conserved structural folds, enabling the discovery of an Acb1 homolog in eukaryotic viruses^64^. With advances in structural predictions and comparative analyses, the field is pivoting towards structure-guided discovery of new anti-defense systems and has been applied to discover new Acrs in phage^65^. However, there is limited representation of small protein structures (i.e. <300 amino acids) derived from phage or MGEs. For context, the Protein Databank (PBD) and AlphaFold Database^66,67^ collectively represent 34,934 phage protein structures, but the Genbank Database contains 13,487 phage genomes that we estimate may encode over 130,000 small accessory proteins with putative anti-defense functions^68^. The next iteration of AntiDefenseFinder will include a database of experimental and predicted protein structures of all known anti-defense systems. We hope it will enable detection of distantly related homologs and allow us to create new HMM profiles to improve detection. In the future, combining protein structural prediction with machine learning algorithms will open a new frontier of anti-defense system discovery.

Despite these limitations, our quantitative detection and analysis of known anti-defense systems revealed fundamental insights into bacterial and phage biology. We observed that over 80% of detected instances of anti-Pycsar protein 1 (Apyc1) were encoded in non-mobile regions of the bacterial genome. Apyc1 was previously identified in phages and prophages and functioned in the degradation of cyclic mononucleotides (cAMP, cGMP, cCMP, cUMP)^31^. Pycsar defense solely relies on cCMP and cUMP^19^ whereas cAMP and cGMP are involved in housekeeping functions^69,70^. However, our evolutionary and functional analyses suggest that Apyc1 originated in bacteria and then phage co-opted Apyc1 to counteract Pycsar defense. An alternative scenario has been observed with Type III CRISPR-Cas defense: Phage encode a ring nuclease (AcrIII-1) that degrades cA_4_ and inhibits CRISPR effector activity^41^ and then bacteria co-opted this inhibitor (Crn2) to regulate CRISPR^42^. Lastly, the recently identified NAD+ reconstitution pathway 1 (NARP1), which inhibits defense systems metabolizing NAD+, was also found in non-mobile regions of bacteria^37^, aligning with our findings and likely plays housekeeping functions in bacteria. Altogether, the AntiDefenseFinder tool has enabled us to explore the diversity of anti-defense proteins across prokaryotes, phages, and MGEs and we hope that we’ve given others the agency to do the same.

## DATA AVAILABILITY

The Anti-DefenseFinder web service can be found at https://defensefinder.mdmlab.fr/. The command line tool is available on GitHub at https://github.com/mdmparis/defense-finder, and its associate MacSyFinder models are also available on GitHub at https://github.com/mdmparis/defense-finder-models. Code and supplementary information are available on GitHub: https://github.com/mdmparis/antidefensefinder_2024 and on Figshare under the DOI: 10.6084/m9.figshare.26526487.

## SUPPLEMENTARY DATA

Supplementary Data are available at NAR Online.

## ACKNOWLEDGEMENTS

We thank Sukrit Silas from the Bondy-Denomy Lab for his insight into the genomics of phage anti-defense systems. We thank the IT Department of the Institut Pasteur for providing support for the AntiDefenseFinder web service.

## Author contributions

E.H. and F.T. conceived the project, and A.B. and J.B.-D. supervised the project and provided feedback. F.T. and J. C. led the development of the AntiDefenseFinder pipeline and HMM profiles, as well as detection and bioinformatic analyses of anti-defense systems. J.C. and M.J. provided support in developing the pipeline and HMM profiles, R. P. developed the web service. E.H. performed the in-depth sequence and structural analyses of Apyc1 homologs. L.W. and J.R. conducted the protein purification and *in vitro* cleavage and kinetics assays of Apyc1 homologs supervised by Y.F.. E.H. and F.T. wrote the manuscript and created the figures, and all authors provided editing and feedback.

