## Supplementary Figures for "Exploring the diversity of anti-defense systems across prokaryotes, phages, and mobile genetic elements"

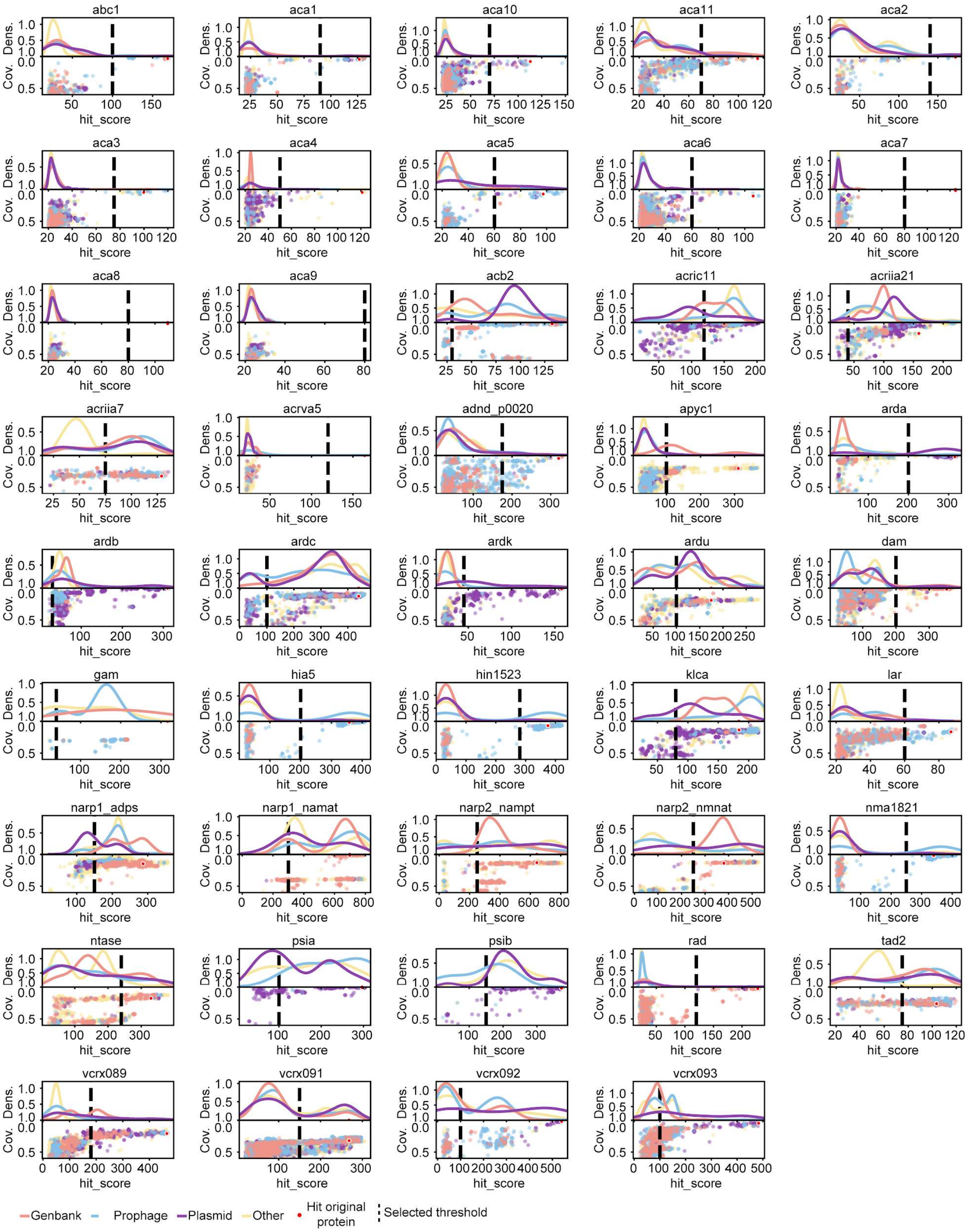


**Figure S1. Anti-defense system threshold choice for developing HMM profiles.** Threshold selection representation for the 44 HMM profiles with at least 1,000 hits in the two databases. Hit score density based on hit localization (top) and distribution of hit score and profile coverage for all hits (bottom). Hits are colored based on their localization (red: phage, blue: prophage, purple: plasmid and yellow chromosome). The selection threshold for each anti-defense protein was manually analyzed and chosen based on the distribution of hits relative to the originally discovered protein. (Plots for all HMM profiles are available on GitHub).

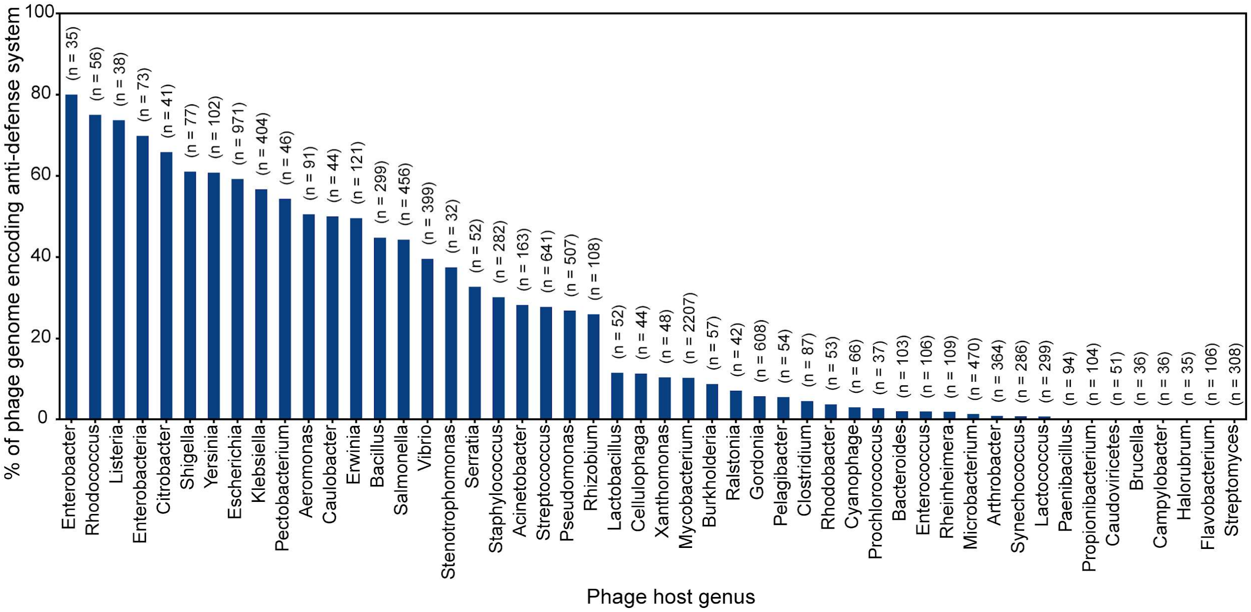


**Figure S2: Phage anti-defense genes are variable across hosts.**

The percentage of phage anti-defense genes categorized by distinct bacteria host genera. In highly studied host genera, the percentage of genome encoding anti-defense genes represent more than 50% of sequenced genomes. In species with less studies on anti-defense proteins, the percentage is much lower (0% for *Streptomyces*), suggesting that a diversity of anti-defense genes may be found in those genera.

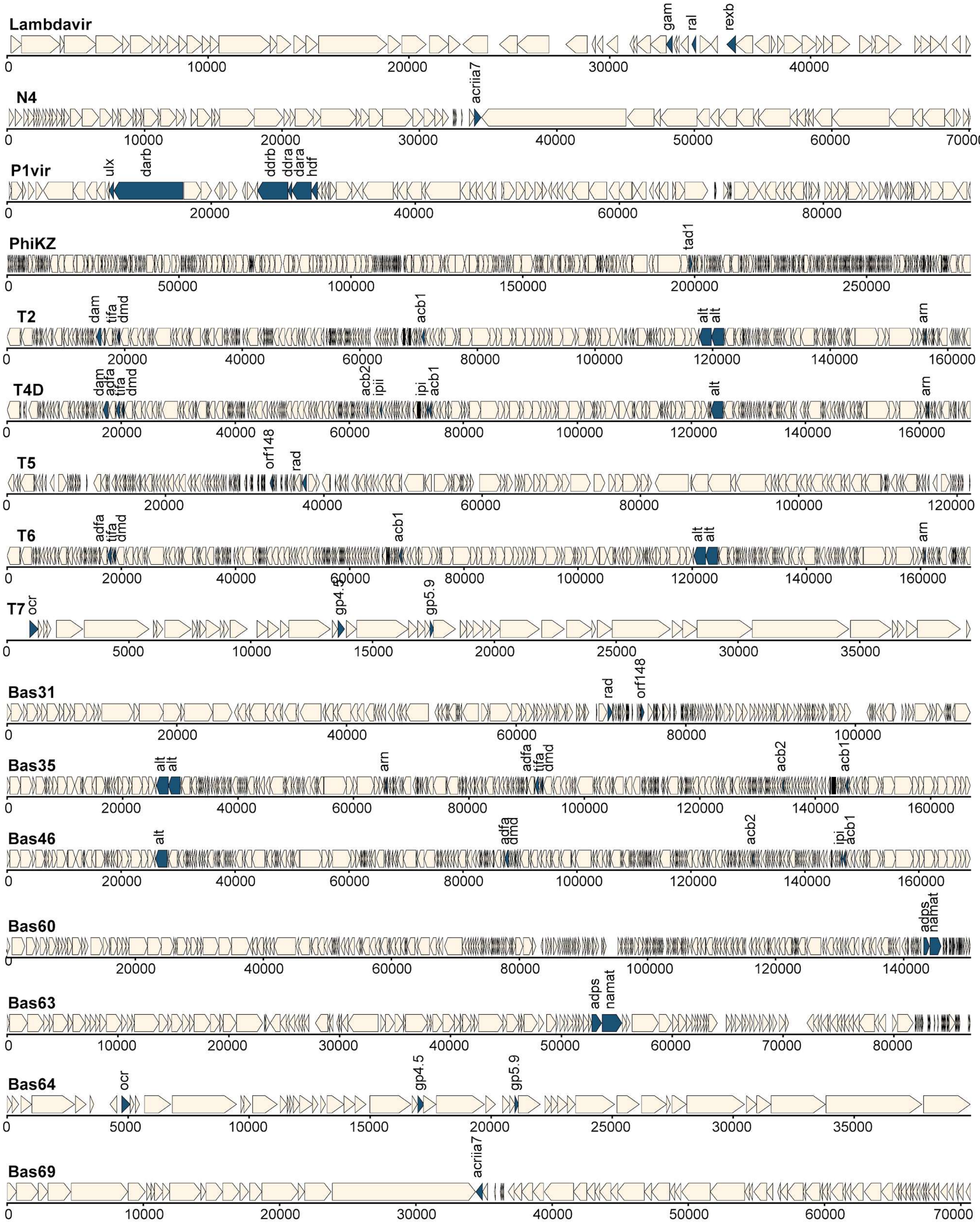


**Figure S3: Phage anti-defense often genes co-localize in genomic loci.**

Example of phage genome encoding anti-defense proteins and co-localization from model phages, as well as phages from the BASEL collection. All detected anti-defense genes are annotated in blue.

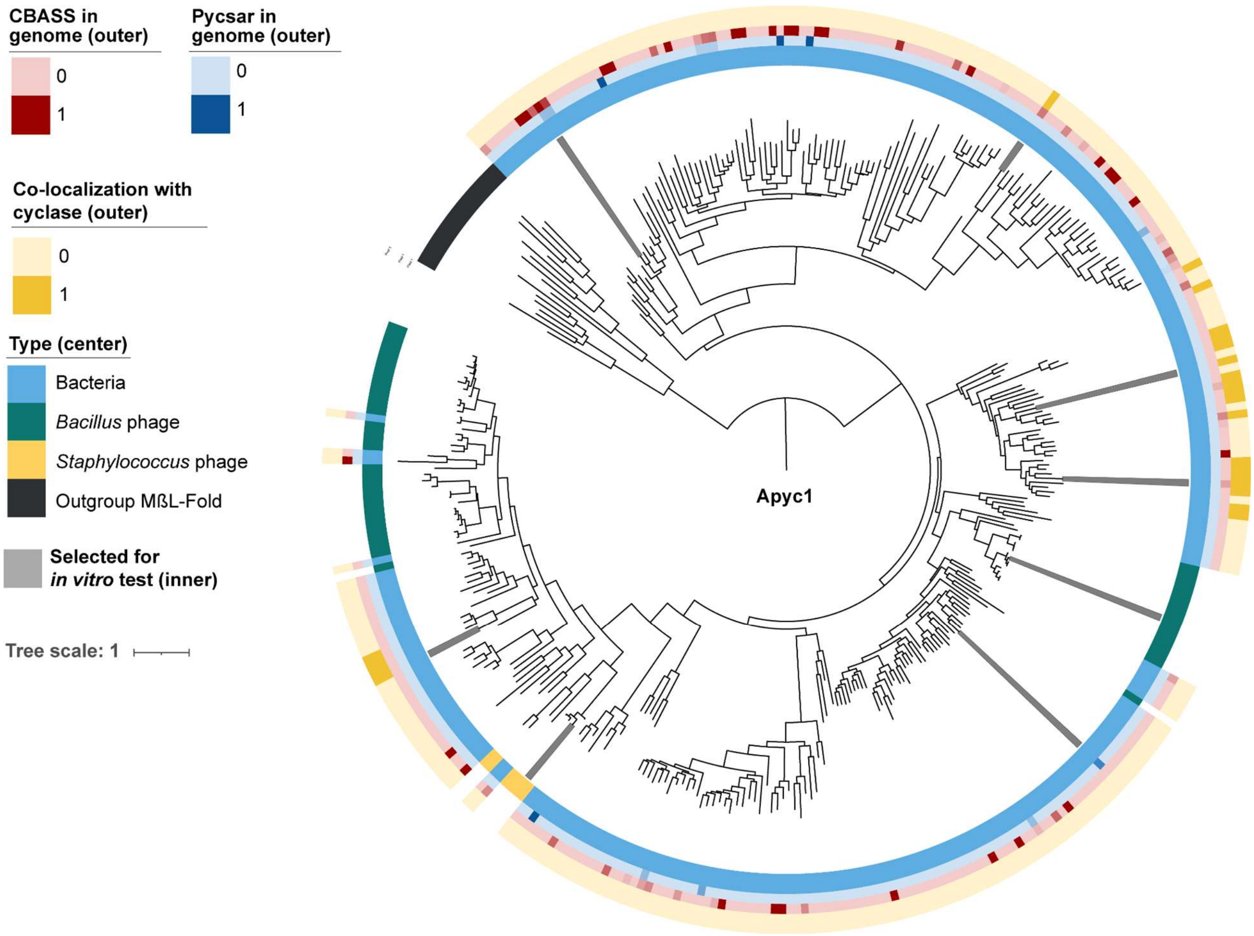


**Figure S4: Apyc1 is encoded in bacterial genomes that harbor cyclases.**
Phylogenetic tree of 356 representative sequences of Apyc1 homologs from bacteria and phage. The outer circles represent the proportion of co-localization with cyclases or the presence of defense systems in the bacterial genome(s) that encode the Apyc1 homolog. The center circle represents the types of Apyc1 homologs: Bacterial genome, *Bacillus* or *Staphylococcus* phage, MβL-fold phosphodiesterases outgroup. Gray highlights the Apyc1 homologs tested for *in vitro* cleavage of cyclic mononucleotides (cNMPs).

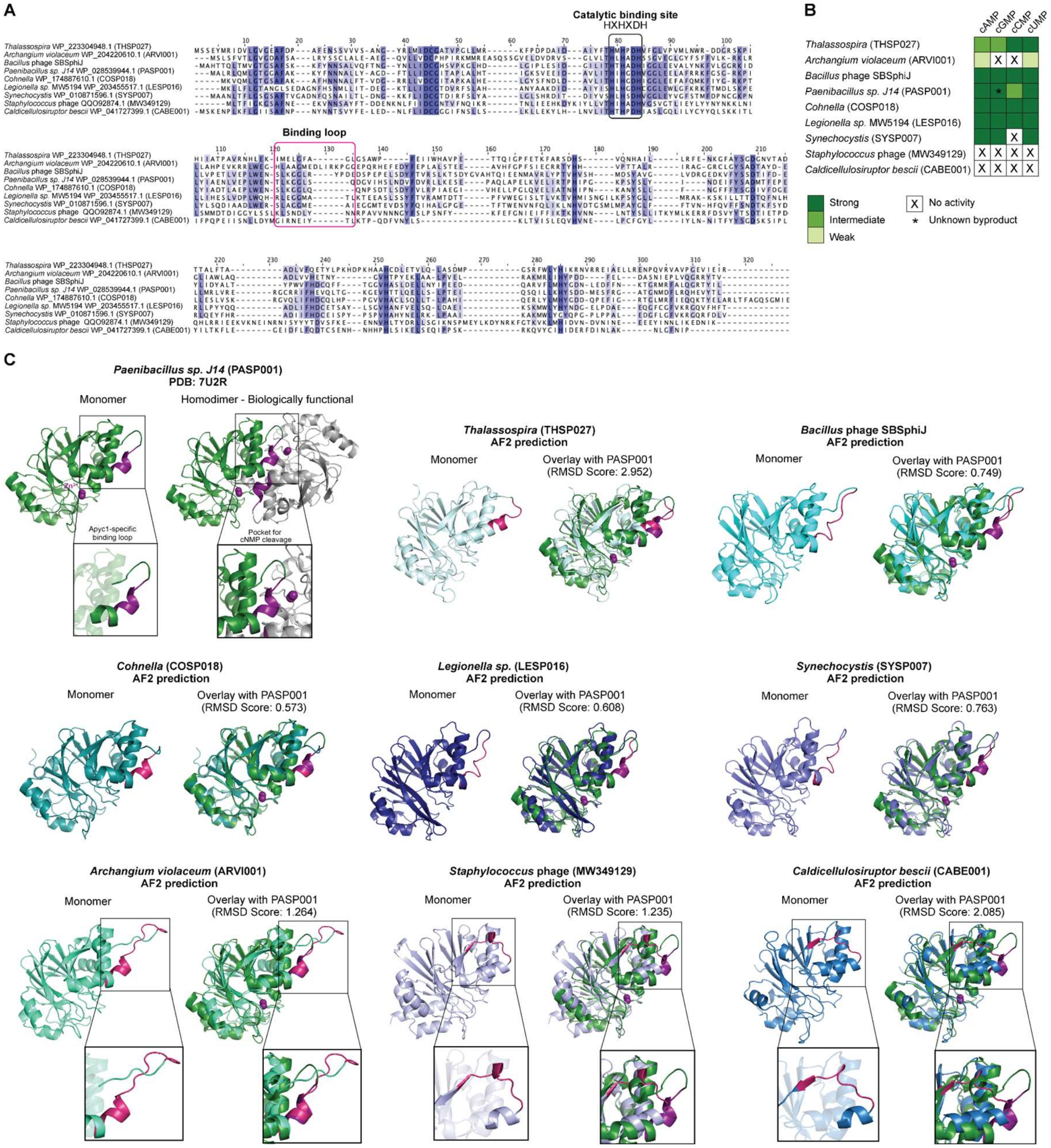


**Figure S5. Sequential and structural differences in Apyc1-specific binding loop.**
**(A)** Multiple sequence alignment of Apyc1 homologs. Residues that are >80 % conserved, >60 % conserved and >40% conserved are shaded in dark purple, light purple, and light gray, respectively. Residues involved in catalysis and binding are circled in black and pink, respectively. **(B)** Summary of the *in vitro* cleavage assay. **(C)** Structures of Apyc1 homologs used in (B). *Paenibacillus sp. J14* (PASP001) Apyc1 was experimentally solved and deposited on the RCSB Protein Data Bank (PBD: 7U2R) as a monomeric unit and homodimeric biological assembly. The remaining Apyc1 homologs were computationally predicted using AlphaFold2 (AF2), and then computationally overlaid with PASP001 Apyc1 using Pymol. Root mean squared deviation (RMSD) scores are provided. Zoomed-in images highlight the Apyc1-specific loop that extends into the cNMP catalytic binding site of the interacting monomer. $Zn^{2+}$ ions that coordinate degradation of cNMPs within the catalytic binding site, as well as the Apyc1-specific loop, are highlighted in purple or pink.

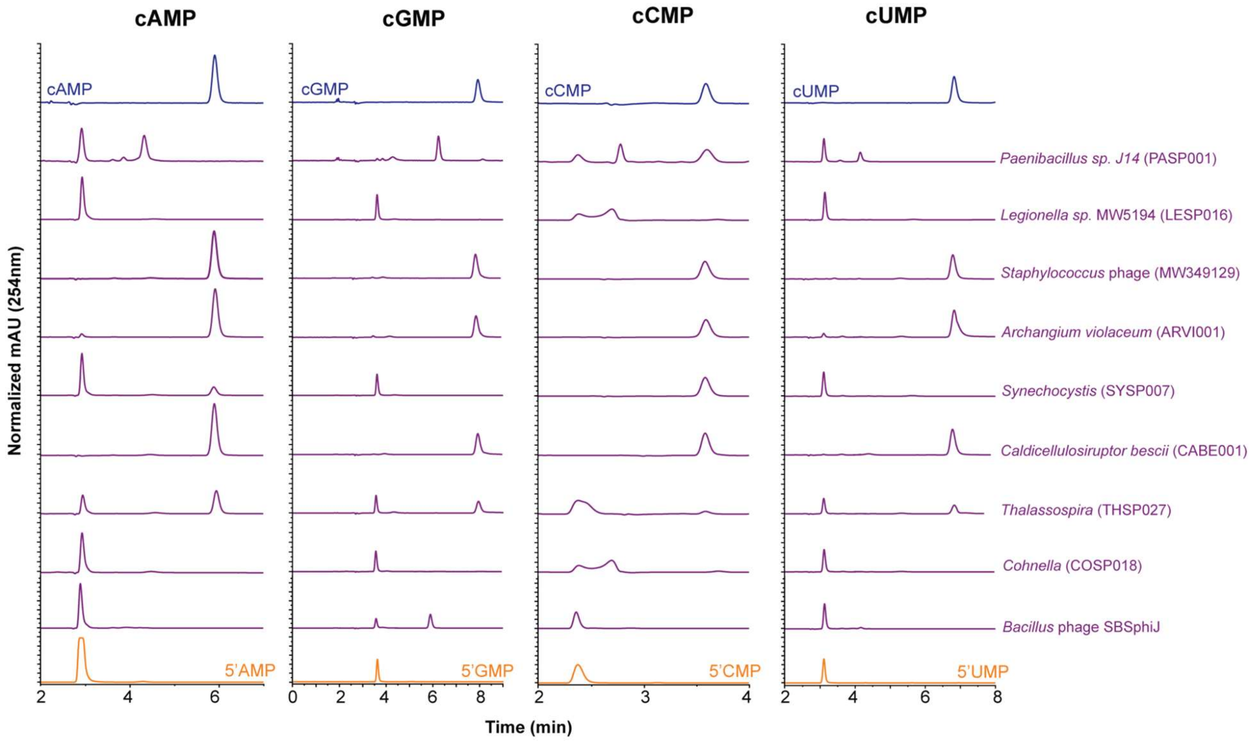


**Figure S6. Apyc1 homologs exhibit a range of cleavage activity.**
HPLC analysis of Apyc1 homologs incubated with the indicated substrates (100 µM) for 20 min at 37°C (n=3).

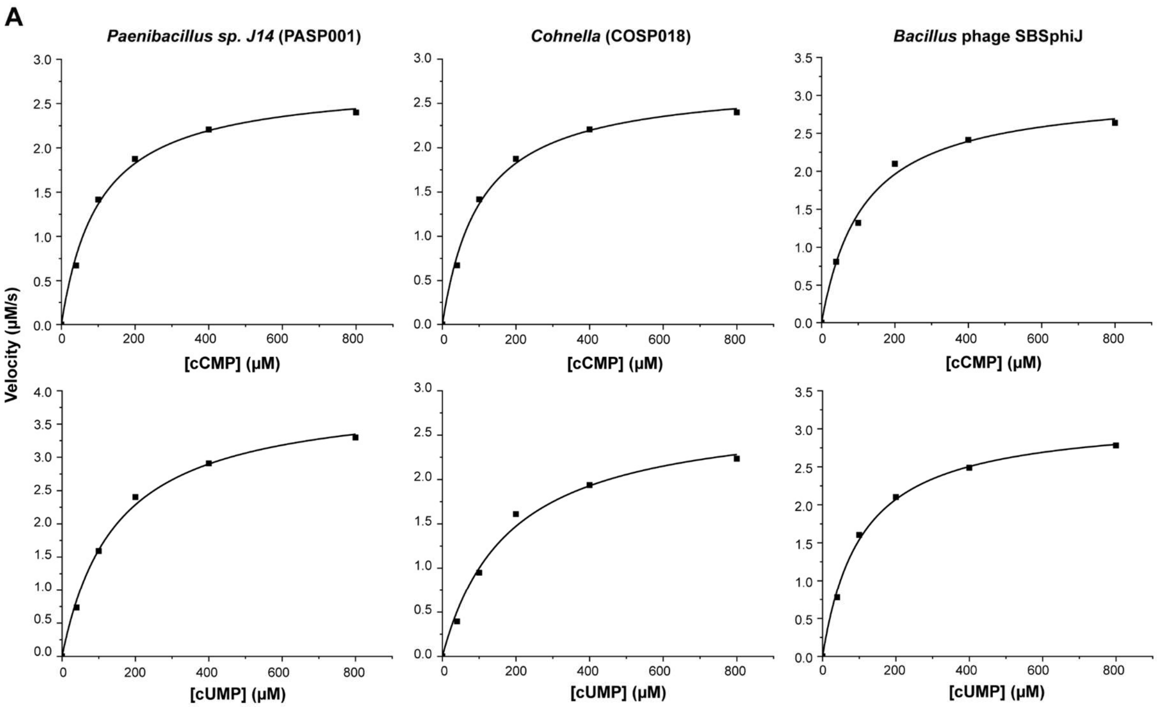


**B**

| | cCMP | | cUMP | |
|---|---|---|---|---|
| | $K_m$ (μM) | $K_{cat}$ ($s^{-1}$) | $K_m$ (μM) | $K_{cat}$ ($s^{-1}$) |
| ***Paenibacillus sp. J14*** **(PASP001)** | 101.51 | 275.18 | 146.18 | 791.79 |
| ***Cohnella*** **(COSP018)** | 101.51 | 1375.89 | 179.34 | 1396.86 |
| ***Bacillus*** **phage SBSphiJ** | 113.17 | 1536.09 | 105.24 | 1581.21 |

**Figure S7. Cleavage kinetics of bacteria and phage Apyc1 homologs.**
**(A)** Kinetic comparison of cCMP and cUMP cleavage by *Paenibacillus sp. J14* (PASP001), *Cohenalla* (COSP018), and *Bacillus* phage SBSphiJ Apyc1 homologs (n=3). **(B)** Summary of kinetic results.
